# Eukan: a fully automated nuclear genome annotation pipeline for less studied and divergent eukaryotes

**DOI:** 10.1101/2025.08.13.670088

**Authors:** Matt Sarrasin, Gertraud Burger, B. Franz Lang

## Abstract

Here, we introduce a new annotation pipeline, called Eukan, designed to deliver reliably high-quality results across a broad range of eukaryotes. First, experimental evidence is automatically leveraged to refine predictions, specifically, RNA-Seq coverage to inform gHMM gene prediction, and intron lengths to inform protein sequence alignments. Second, a consensus is created from an empirically optimized weighting of gene models from multiple sources. Third, Eukan runs a post-annotation routine to recover gene models missing from the consensus that otherwise have strong transcript support and appear to be protein-coding. We compare the results of Eukan with those of three popular freely-available pipelines (Maker, Braker, Gemoma) on 17 phylogenetically diverse haploid and diploid nuclear genomes. In addition to the commonly reported annotation accuracy statistics, we define a novel classification system of critical defects commonly observed in automated annotations. Furthermore, we developed a statistical model that demonstrates each of the tested pipelines correctly identified the majority of the validated “Gold Standard” gene models across the test set, but each pipeline uniquely generates a non-negligible portion of either fragmented, artificially fused, or missing gene models. Despite that, we demonstrate that Eukan performs consistently well where other pipelines encounter challenges, such as for compact protist genomes.

**Contact:** Matt Sarrasin;

## Introduction

Genome sequencing projects typically involve three principal, successive steps, first, genome assembly, second finding genes within that assembly (gene identification) and prediction of their structures (exons, introns, and UTRs), and third, functional annotation. Automated methods in gene prediction (also known as ‘structural genome annotation’, or simply ‘gene annotation’ as referred to herein) have not kept pace with the ever accelerating rate of data generation, particularly for eukaryotes. The reason is the diversity and complexity of nuclear genome architectures, which continue to pose challenges to automation. Such challenges even apply to well-studied model eukaryotes where manual expert curation regularly constitutes a large part of gene modelling efforts (Salzberg, 2019).

The conceptual framework for annotation as reviewed elsewhere (2) is still relevant today. The core components include (i) pairwise alignments (via, e.g., Blast (3), Exonerate (4), etc.) of organism-specific transcript sequences reconstructed from RNA-Seq experiments if available, and/or pairwise alignments of amino acid sequences of phylogenetically related organisms to the target genome; (ii) gene prediction algorithms, such as generalized Hidden Markov Models (HMMs), trained on RNA-Seq read mapping (via Hisat (5), Star (6), etc.) or sequence alignments; (iii) evidence integration and transcript isoform modelling to create consensus models from the first two steps; or (iv) some combination of the three components above.

Yet, at the time of writing, numerous challenges remain. Annotation pipelines strive to replicate the complex, multi-step decision-making processes of an expert curator (Brent, 2008). These curators employ various analytical tools to either make gene-specific assessments or to come to a decision with a degree of uncertainty, both of which are difficult to automate. In addition, while human curators are typically experts in the biology of a given organismal group, automated annotation must be a ‘one size fits all’ procedure for datasets from a broad range of eukaryotic taxa whose genomic features can differ substantially and often contain biases that lead to misinterpretation. In particular, the quality of the underlying genome assembly bears significant weight in determining the quality of annotation (7, 8), but this subject is beyond the scope of this study.

It should be noted that existing annotation pipelines are mostly focused on identifying and modelling protein-coding genes, while genes specifying structural and regulatory RNAs (ncRNAs) are largely neglected. One reason for that is that many classes of ncRNAs remain poorly understood, or are poorly conserved between lineages and therefore difficult to identify (9). The few ncRNAs that can be predicted with confidence by their conserved sequence and secondary structures include tRNAs, rRNAs, spliceosomal RNAs, RNase P RNA and MRP RNA (10). For a complete list see the current version of RFAM.

### Strengths and limitations of current automated eukaryotic gene annotation pipelines: Maker, Braker, and Gemoma

**Maker** (11) was the first pipeline to synthesize coding gene models informed by evidence derived from a variety of raw input files: the genome assembly itself, nucleotide sequences from a transcriptome assembly, and protein sequences from related species. Maker’s conceptual design is centred around integrating various forms of evidence and automating steps that were previously difficult to achieve. First, Maker will automatically align transcript and protein sequences (specified by the user) to the genome assembly to generate preliminary gene models. Second, the user extracts structural information from those models, with the help of auxiliary scripts, to train a variety of gene prediction algorithms including Augustus (12). Maker then runs automatically the trained gene predictors to generate *ab initio* gene models. Subsequently, the user must run Maker again to automatically select and modify *ab initio* gene models that are most congruent with protein and transcript sequence alignments, with the option to emit predictions without sequence alignment support (11). Despite those advancements, the user is still required to manually intervene at several points, which requires knowledge about parameterization related to the various manually-driven tools. Maker continues to be maintained but no fundamental changes have been made to its design since its second major version (13). As argued by Hoff et al (2016), an issue with Maker’s approach is that it suffers from errors such as mis-assemblies and fragmentation introduced by transcript reconstruction algorithms.

To overcome such pitfalls, **Braker** was developed, which leverages unassembled RNA-Seq read coverage at exon-intron junctions (i.e., split reads). In addition, the Braker pipeline is fully automated in a single script, thus reducing interventions by the user. Braker’s parameterization is tailored to the type and availability of data at the user’s disposal, e.g., options to derive hints from RNA-Seq and/or homology data (inferred from protein sequence alignments), and options specifically for fungal or grass genomes. The final gene models are generated by the single and (in theory) well-informed gene prediction algorithm, Augustus (14). However, relying on decisions made by a single predictive algorithm about gene structure is a double-edged knife, since accuracy is heavily dependent on parameterization as well as the number, representativeness and diversity of both gene models for training and external evidence given to Augustus. Recently, Braker has been upgraded (version 2) to optionally make use of protein sequence alignments as evidence of coding DNA sequence (CDS) (15).

Around the same time Braker (version 1) was released, **Gemoma** was published, a fully automated annotation package (16). Version 1.1.1 foregoes GHMM-based gene prediction algorithms altogether. Instead, this pipeline exploits the conservation of intron positions (17) identified by protein sequence alignments for building gene models. The most recent version of Gemoma makes use of intronic coordinates inferred from RNA-Seq in conjunction with those inferred from sequence alignments of proteins from other species (19). While homologous sequences potentially provide high-quality support to a gene model, the risk is twofold. Not only will utility drop as a function of evolutionary distances of the organism of interest, but also errors within those homologous sequences provoke missing, fragmented and incorrect gene models (20).

The three pipelines discussed above are considered to be applicable to eukaryotes in general, however, little is known about their performance beyond a handful of well-studied animal, plant and fungal genomes on which they have been tested. To our knowledge, no comprehensive performance report on any of the numerous other and most diverse eukaryotic lineages, collectively and informally referred to as ‘protists’, have been published.

Here we present a new pipeline, Eukan, that exploits more information than traditional annotator software. Benchmarking was performed against Maker, Breaker, and Genoma, by measuring performance against a set of genomes that is larger and taxonomically much broader than usually done, including not just the plant *Arabidopsis thaliana* and the two model animals, *Caenorhabdhitis elegans* and *Drosophila melanogaster*, but also Apicomplexa, Chlorophyta, Fungi, Kinetoplastida, and Rhodophyta.

## Methods

### Eukan workflow

Figure 1 depicts an overview of Eukan. The pipeline minimally requires a genome assembly (in fasta format) in which repetitive regions were soft-masked (e.g., with RepeatMasker (21)), along with at least one complete proteome from an organism as phylogenetically close as possible (sequences of amino acids in fasta format). Additional proteomes from more or less closely related organisms improve the results. For optimal runs, additional input is required, for example, transcript sequences assembled from RNA-Seq reads (e.g., using Trinity (22)) and aligned to the genomic contigs (e.g., with STAR (6)). The other consists of RNA-Seq read coverage of the genome (extracted from the STAR BAM file; see the Pre-processing Section below for details). The third additional input describes intronic coordinates inferred from split RNA-Seq reads mapped to the genome (extracted from STAR output). All three input are provided in the general feature format (GFF) and the corresponding files are referred to in the following as transcript, RNA-Seq-read coverage, and intron coordinate files.

**Figure 1.**
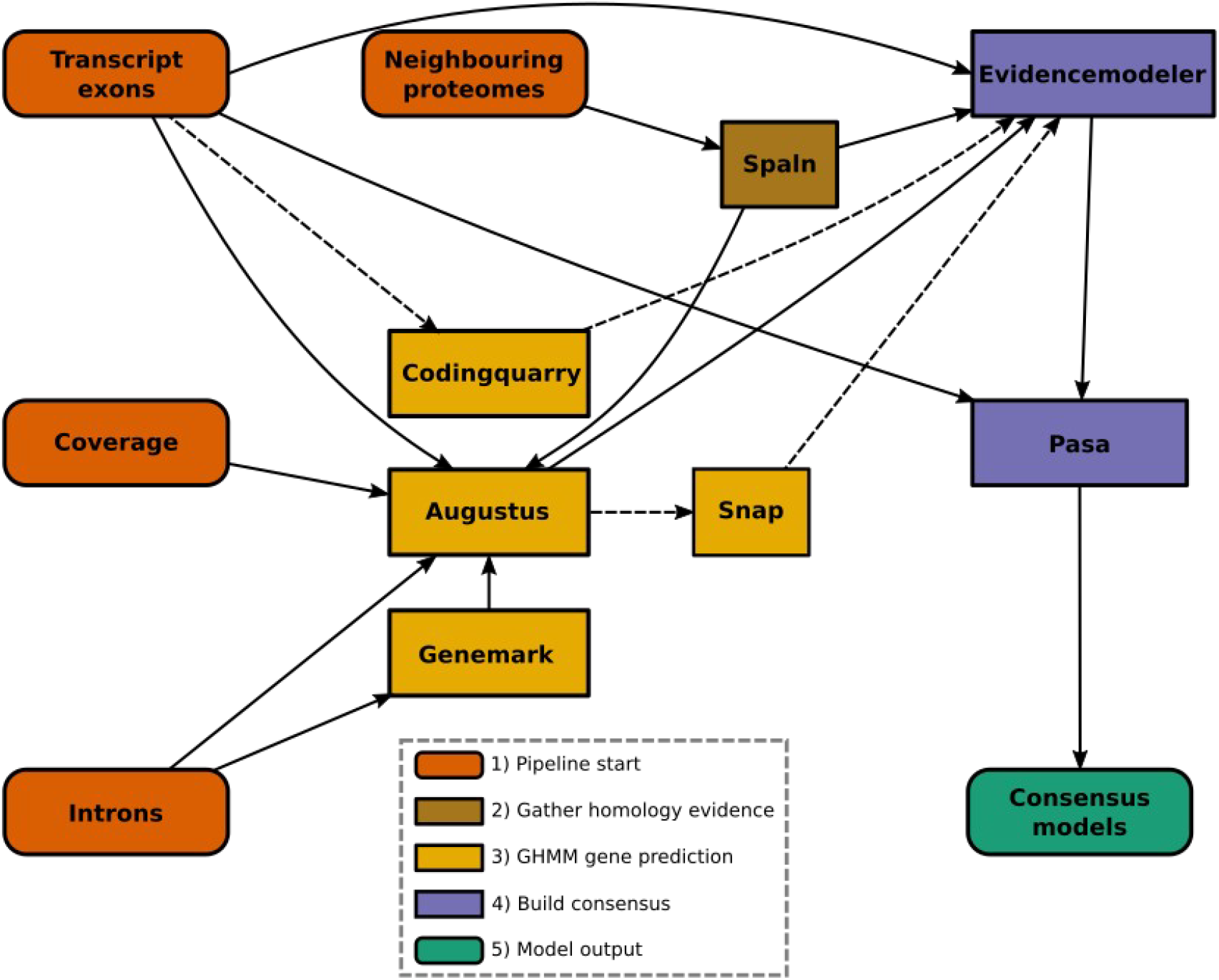
Flowchart of the steps involved in Eukan. Initially, exon features of transcripts (in GFF format) provided to CodingQuarry if the target organism is a fungus. ORF coordinates are also extracted from exons to provide to Augustus at a later step. Next, protein sequences are aligned to the genome using Spaln, followed by initial gene predictions by Genemark. Concordant gene models between Genemark predictions, transcript ORFs, and protein sequence alignments are identifed and used as a training set for Augustus. If the target organism is a fungus, CodingQuarry is run, and Snap trained on the Augustus models with the maximum score of 1. Aligned transcripts, protein sequences and gene predictions are combined into a single consensus using EvidenceModeler with weights 3, 2 and 1, respectively. Finally, consensus models are handed over to Pasa to model alternative splicing according to the input transcript sequences, and UTRs are added to predicted transcripts.

The initial steps of Eukan involve building a collection of preliminary gene models for the purpose of training the Augustus gene predictor (12). First, the CDS regions of transcripts are identified. Next, the intron length distribution is inferred from the intron coordinates using fitild (23). This information will instruct the alignment of the protein sequences from the species of interest with those from close relatives (performed by Spaln (24); yields a GFF file of alignments). Finally, gene models are predicted with Genemark (25) run in ‘ET’ mode whereby intron coordinate information is provided.

To train Augustus, we use the inferred gene models whose structures are concordant between at least two out of the three sources: transcript-contained open reading frames (ORFs), protein sequence alignments and Genemark predictions. After training (optimized using the software’s helper script), Augustus is then run on the genome assembly providing supporting information commonly referred to as ‘hints’ (typically in GFF format). As required by Augustus, the technical terms specifying these hints are ‘exon’ from the transcript file, ‘intron’ and ‘exonpart’ denoting coverage from the RNA-Seq-read coverage file, and ‘CDSpart’ hints from inter-species multiple protein sequence alignments.

Two additional gene prediction algorithms are executed in the case that the target genome is specified as fungus via a command line option (−-fungus). First, Codingquarry (26) is called whereby transcript-genome alignments are provided as input. Second, Snap (27) is also called, initially trained on Augustus predictions that have a score of 1.

In the last steps, the GFF files from transcript alignments, protein alignments, and Augustus gene models (plus Codingquarry and Snap models if available) are given to Evidencemodeler (28) to create consensus gene structures according to the following weighting scheme: weights of 1, 2 and 3 are assigned to gene predictions, protein sequence alignments, and transcript alignments, respectively (empirically determined, data not shown). To model alternative transcripts and add corresponding UTRs, the consensus models generated by Evidencemodeler are provided to Pasa (28), along with the transcript file.

The final (optional) step of the pipeline is to assign functional information to the gene models. Conceptually translated sequences of transcript models are searched against the Swiss-Prot reviewed protein collection (https://ftp.uniprot.org/pub/databases/uniprot/knowledgebase/complete/) using phmmer, as well as searched against the Pfam database (http://ftp.ebi.ac.uk/pub/databases/Pfam/releases/) using hmmscan (29). The best non-overlapping hits below a specified e-value threshold provide information about validated, expert-curated homologs of a given gene model and conserved protein domains. All this information is documented in the genome-annotation GFF3 file, which, for conveniance, is formatted such that it can be processed directly with, for example, NCBI’s submission tools without any further interventions.

### Classification of gene prediction outcomes

We built upon the statistical framework described above by extending the prediction classification at the level of genes, transcripts, exon (strictly the coding coordinates), and introns. Our classification framework defines terms similar to those implemented in the Busco tool (e.g., ‘complete’, ‘fragmented’, ‘missing’). The difference is that Busco classifications are defined at the protein sequence level, whereas the classes defined here are based on the overlap of genomic feature coordinates.

A gene identified at a given locus is categorized as exact, inexact, missing, merged or fragmented (Figure 2). Further, a prediction is deemed a ‘match’ if its genomic coordinates overlap with those of a single reference, either coinciding exactly (no false positive or false negative loci) or inexactly (some degree of false positive or false negative loci) at the 5’ and 3’boundaries. In contrast, an incorrect prediction, or ‘defective’ model, is either ‘merged’, ‘missing’ (i.e., FN) or ‘fragmented’. The coordinates of an artificially ‘merged’ gene prediction span those of two or more references (by more than 50% of the reference lengths in question). This typically occurs when an intergenic region is mistaken as an intron. A ‘missing’, or FN, prediction occurs when a reference gene is mis-identified as an intergenic region. Finally, two or more distinct gene predictions that each overlap (by at least 50%) separate portions of a single reference gene are defined as ‘fragmented’. This typically occurs when an intron is mistaken as an intergenic region.

**Figure 2.**
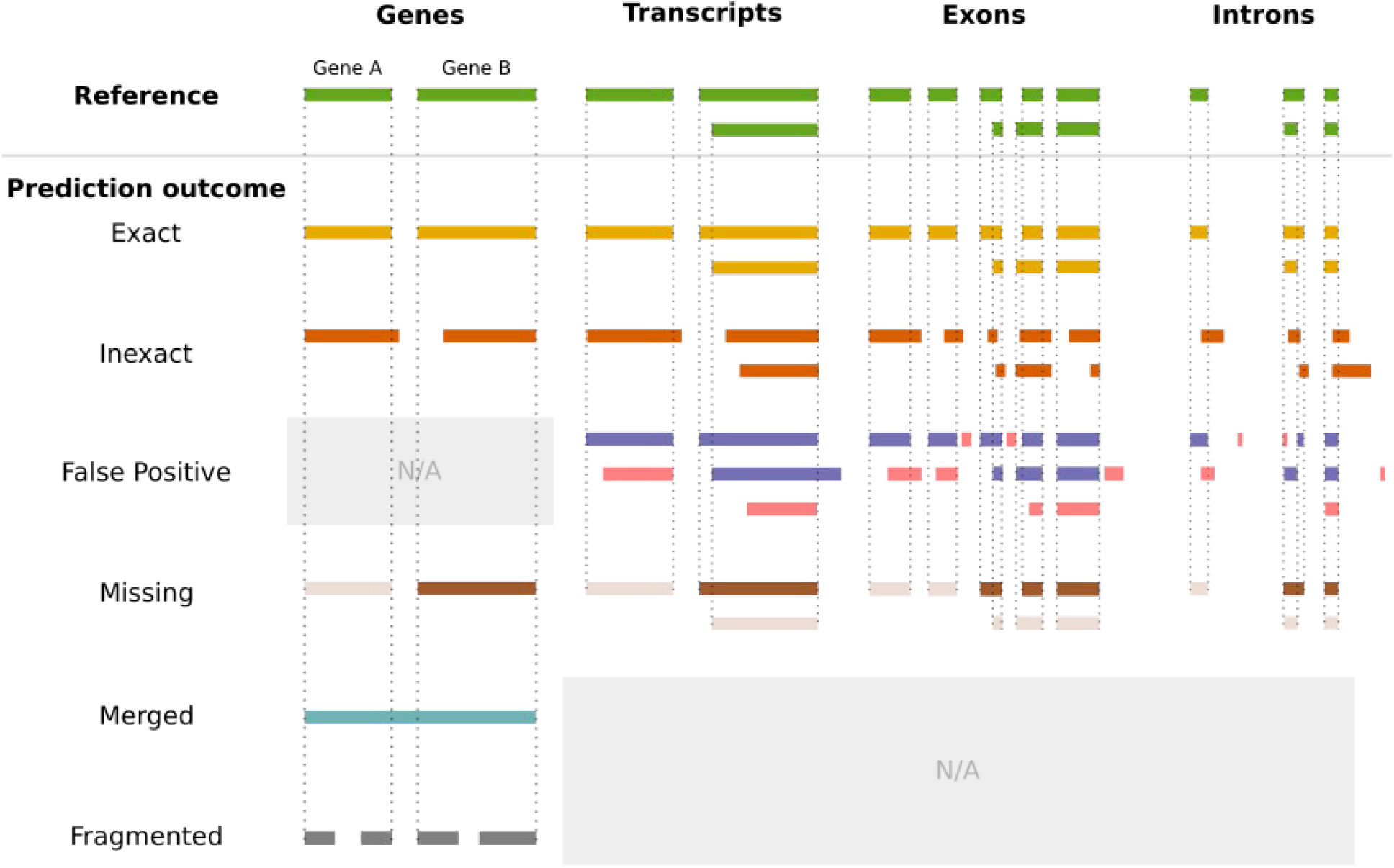
Outcomes of gene predictions compared to the (true) gene structure (reference) as defined in this work. Prediction coordinates can be either an exact (perfect) match or an inexact (imperfect) match with the reference; completely missing (bars in light brown); a merge of two or more distinct, adjacent models; or a fragmentation (split) of a distinct gene into multiple components. False positives are strictly defined for transcripts, exons and introns (not genes, hence the gray box), and are represented in the pink bars. ‘Merged’ and ‘fragmented’ are strictly defined for gene models, hence the grayed out area for transcripts, exons and introns.

At the transcript level, predictions are classified as either a ‘match’, ‘missing’ or ‘false positive’ (FP). To account for multiple transcript models per gene, ‘matching’ transcripts are identified by computing the maximum pairwise sum of nucleotide overlaps between prediction and reference. Reference transcripts with lacking prediction are counted as ‘missing’, whereas superfluous transcript predictions are counted as false positives.

Intron and exon predictions are grouped by their parent transcript in order to compute maximum pairwise overlaps with their respective reference coordinates. Introns and exons with maximum pairwise reference overlap length are selected as ‘matching’ predictions. Like missing and false transcripts, lacking exon and intron predictions are counted as ‘missing’, and superfluous predictions are counted as false positives.

## Results & Discussion

### Beyond accuracy statistics for pipeline performance assessment

The accuracy statistics Sn, Pr and F1 remain the primary methods for benchmarking pipelines. One study has also made use of the Chi-squared test based on those statistics (30). However, the disadvantage with these aforementioned methods is that they are inherently diagnostic, which implies that they reveal little about the predictive power and shortcomings of a pipeline. Performance assessments have been restricted to these statistics in the past due to sparse availability of high-quality, verified structural information about genes. Another consequence of this lack of data availability is that a narrow set of organisms *(Arabidopsis, Caenorhabditis, Drosophila* and fission yeast) have been *de facto* subjects for annotation performance comparisons (13, 14, 19). Such a necessarily limited scope leads to biases in the performance of a pipeline. Only recently, studies are including additional organisms beyond the usual three or four suspects, such as certain emerging model fungi (*Verticillium dahliae* and *Plicaturopsis crispa (30)*), plants (*M. truncatula, S. lycopersicum*) and animals (*X. tropicalis, D. rerio, B. terrestris*) (15), as well as phylogenetically close neighbours to existing model organisms, e.g., *Caenorhabditis (19)*. Still, taxa outside fungi, plants and animals (i.e., protists) remain largely absent from comparisons (31) despite the fact that members of this group represent the majority of eukaryotic genomic diversity (32).

To our knowledge, the 17 organisms selected in this study represents the most diverse and comprehensive test set reported on pipeline comparisons to date. Such a test set permits statistical analyses that provide some notion of generalizability, as well as reveal patterns of behaviour in pipelines that were previously unattainable with fewer organisms. Specifically, a relationship can be inferred in feature predictions (genes, transcripts, exons, introns) found to be correct or incorrect with respect to the corresponding number of reference features. Further, cumulative empirical distributions of F1 scores computed for correctly predicted features can be generated in order to test hypotheses of significant differences in accuracy between pipelines. Lastly, it becomes possible to test for differences in whether pipelines tend to infer the same coding regions, as well as to test whether predictions from various pipelines at a given genomic locus tend to match the reference.

### Extending the curated reference model collection by Busco analyses

As a ‘Gold Standard’ for pipeline performance assessment, we used curated (SwissProt) models of protein-coding genes where available (Supplementary materials) specifically for the reference genome accessions listed in Methods. While *O. sativa, S. pombe, S. cerevisiae* and *D. melanogaster* had more than 2000 curated models and *A. thaliana, C. elegans* and *N. crassa* several hundred, only a handful were available for e.g., *Aspergillus* and *Ustilago* (Supplementary Table 1). We reckoned that this bias could be counteracted, at least in part, by supplementing curated models with uncurated models reported to be complete by the tool Benchmarking Universal Single-Copy Orthologs (Busco). The hypothesis is that the quality of uncurated models found to be ‘complete’ (by Busco) should be similar to that of curated models found to be complete, and thus can be reasonably considered ad hoc Gold Standard models.

To test this hypothesis, a three-fold approach was implemented. First, Busco was run on the conceptually translated sequences of all reference models for each species using the closest available OrthoDBv10 lineage dataset to the species. The result of this assessment suggests that the ‘completeness’ rate for the 17 organisms was found to be fairly high, at 91.4% on average and a 97.1% median. The average rates of ‘missing’ and ‘fragmented’ Busco genes were thus found to be correspondingly low, at 7.0% and 1.5% respectively, with median rates of 1.6% and 0.80% respectively. Second, all reference models identified by Busco (‘fragmented’ and ‘complete’, according to its evaluation scheme (33)) were partitioned into one of two groups, either SwissProt-curated or uncurated. A two-sample z-test of proportions of fragmented-to-complete between these two latter groups revealed no significant difference, which suggests that the outcome of Busco assessments in uncurated models is similar to that of curated models. In a third step to provide better visibility into the underlying quality, the cumulative F1 distributions were collected from gene predictions (by Braker, Eukan, Gemoma and Maker) against both uncurated reference models identified as ‘complete’ by Busco, and SwissProt-curated models, for all species (Supplementary Figure 3). A two-sample test, based on the DTS test statistic, revealed no significant difference between F1 distributions of curated vs uncurated models, which suggests that the expected quality of a gene prediction against a curated and an uncurated (but ‘complete’ as determined by Busco) model is indistinguishable. Taken together, these results suggest that the quality of reference models identified as ‘complete’ by Busco is reasonably similar to that of SwissProt curated models and can therefore serve as a complement.

Supplementary Table 1 shows the breakdown of the extended gold standard model counts. ‘Curated non-Busco count’ are SwissProt curated models not within the Busco set, whereas ‘Busco uncurated models’ were found to be complete by Busco but remain uncurated. The intersection of the SwissProt and complete Busco models are shown under ‘Curated and Busco gene overlap’. In sum, the number of gold standard reference models is increased substantially when including those found to be complete by Busco. Even the gold standard sets for well known model organisms like *S. pombe* (21% increase to 3174), *A. Thaliana* (84% increase to 1886), *D. Melanogaster* (56% increase to 4688) and *C. elegans* (92% increase to 3348) benefit from the inclusion. The benefit is equally visible for other organisms that do not receive as much manual curation effort, increasing the number of gold standard models to hundreds (all protists) or even thousands (*N. crassa, A. nidulans, U. maydis*, and all plants).

### Summary of predictions on the extended gold standard collections

A more detailed description of gene predictions at levels of gene, transcript, exon and intron predictions, are available in the Supplementary Materials. To summarize, assessment of gene predictions generated by the tested pipelines (Braker, Eukan, Gemoma, and Maker) on the extended reference collections revealed generally high rates (>75%) of ‘completeness’, according to Busco standards, where results are more often similar across pipelines. Some of the differences occurred where Maker and Braker exhibited high rates of ‘missing’ models in specific organisms. Despite those differences, all pipelines demonstrated strong correlations in matching gene predictions to reference loci, with most predictions being exact matches. Again, some outliers were noted in some organisms, particularly for Gemoma and Maker. In transcript predictions, discrepancies in duplicated sequences were observed, especially in organisms with known alternative splicing, yet all pipelines generally produced high F1 scores for matching transcripts. For exon and intron predictions, all pipelines successfully identified over 95% of reference exons and introns, with Braker generating more false positives than others, while Eukan consistently showed lower rates of erroneous predictions.

### Gene models assessed by Busco for pipeline comparison

Quality assessments of automated gene building require knowledge about gene structure correctness on a genome-wide scale, but that information is often limited to a handful of expertly verified models, if any at all. Even manually curated models can be imperfect due to misassembly, curation error or uncertainty. Automated Busco analyses can supplement quality assessments by classifying the protein sequences as complete, fragmented or missing, but they do not provide information about the correctness of the gene’s exon/intron structure. Despite these limitations, including reference models deemed ‘complete’ by Busco increases the breadth of organisms and the depth of models available for the purpose of benchmarking pipelines in this study. As supported by the observation described in the Results section, the prediction accuracy (F1) on both curated and non-curated, Busco-complete reference models are similar for most pipelines. In fact, Busco-complete curated and non-curated models were generally predicted accurately by at least three out of four annotation pipelines. Only around 5% of proxy curated models were found to be faulty due to potential Busco misclassification, incorrect reference coordinates (generated by automated procedures), assembly error, or some combination thereof and thus should not have a considerable impact on pipeline benchmarking.

Given that 1) no difference was observed in proportions of ‘complete’ and ‘fragmented’ models, as deteremined by Busco, for curated vs uncurated reference models, and that 2) no statistical difference is observed between cumulative prediction F1 distributions, we are confident to use Busco-complete uncurated reference loci as proxy curated models in order to enhance the set of gold standard models, particularly for organisms with few or no Swiss-Prot entries.

### Gene predictions generally succeed but beware exotic genomic properties

All pipelines identified roughly the same set of reference gene models (Figure 4) with most of the correct gene predictions being exact matches to the corresponding reference coordinates (Supplementary Figure 6). Rigorous performance comparison shows that no statistically significant difference in accuracy was observed between the pipelines, and none of the tested pipelines significantly under-or outperformed globally (Figure 3). Still, Maker, had a significantly higher tendency to predict a single model at a given locus where the reference (and other pipelines) suggest multiple distinct gene models. The minority of predicted genes (∼5%) with some defect are not shared by the pipelines but rather appear to be pipeline-specific (Figure 4). Nevertheless, Eukan was the only pipeline to not exhibit outlier behaviour in gene predictions like what was observed in Maker’s predictions in *O. Sativa, S. cerevisiae, L. major, T. gondii,* Gemoma’s predictions in *C. primus, C. reinhardtii, D. discoideum,* or Braker’s predictions in *S. pombe, C. merolae, T. brucei*.

**Figure 3.**
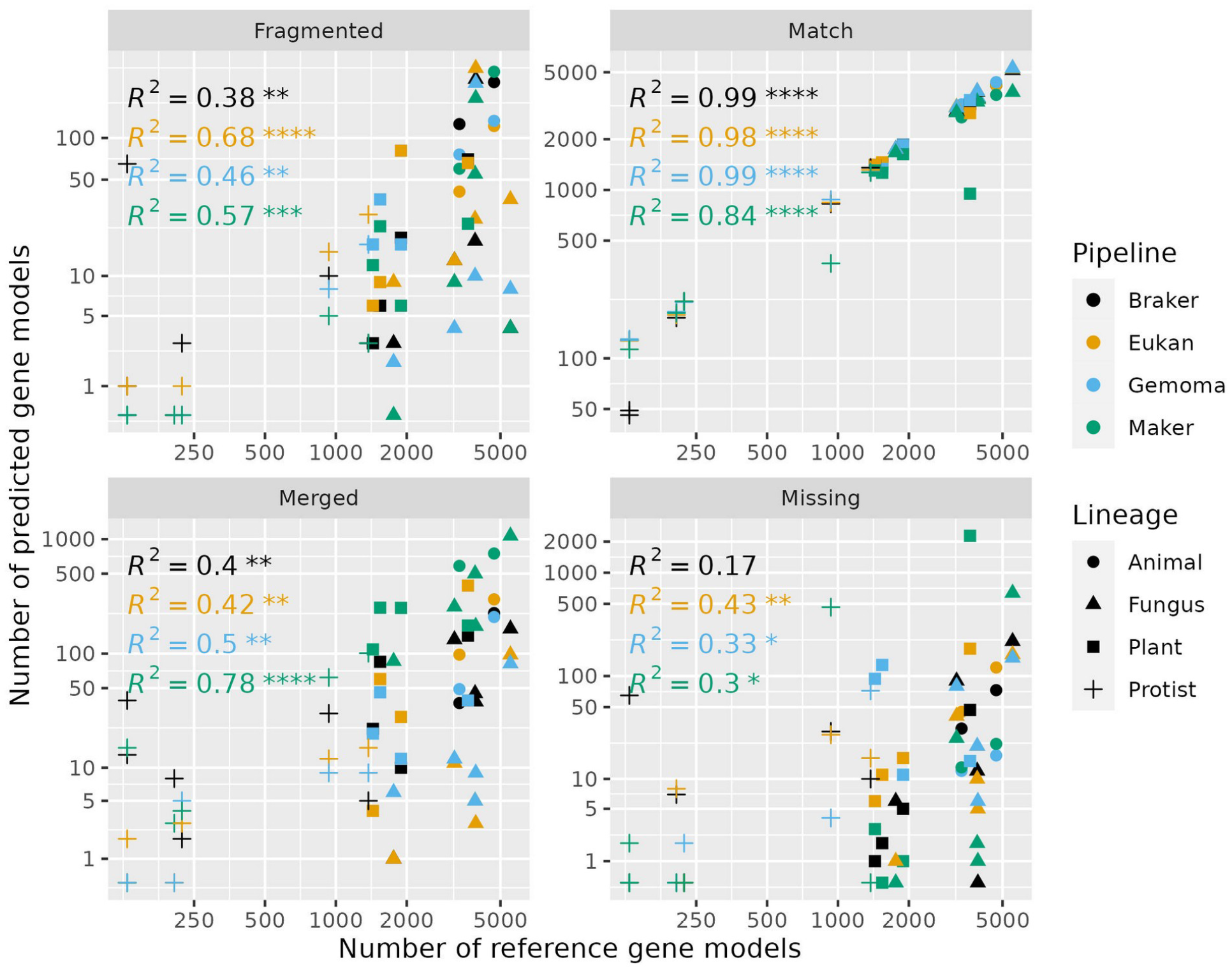
Scatterplots of the number of matching, missing, split and false positive gene predictions as a function of the number of reference genes (curated, Busco-complete, or both) per annotation pipeline for each of the 17 tested genomes. ‘Animal’ includes C. elegans, D. melanogaster; ‘Fungus’ includes A. nidulans, N. crassa, S. cerevisiae, S. pombe, U. maydis; ‘Plant’ includes A. thaliana, O. sativa, C. reinhardtii, C. primus, O. lucimarinus; ‘Protist’ includes P. falciparum, T. brucei, D. discoideum, L. major, C. merolae, T. gondii. R^2^ represents the coefficient of determination, and the associated numbers of asterisks represent the level of significance (^*^ : p-value <0.01; ^**^ : p-value <0.001; ^***^ : p-value <0.0001).

**Figure 4.**
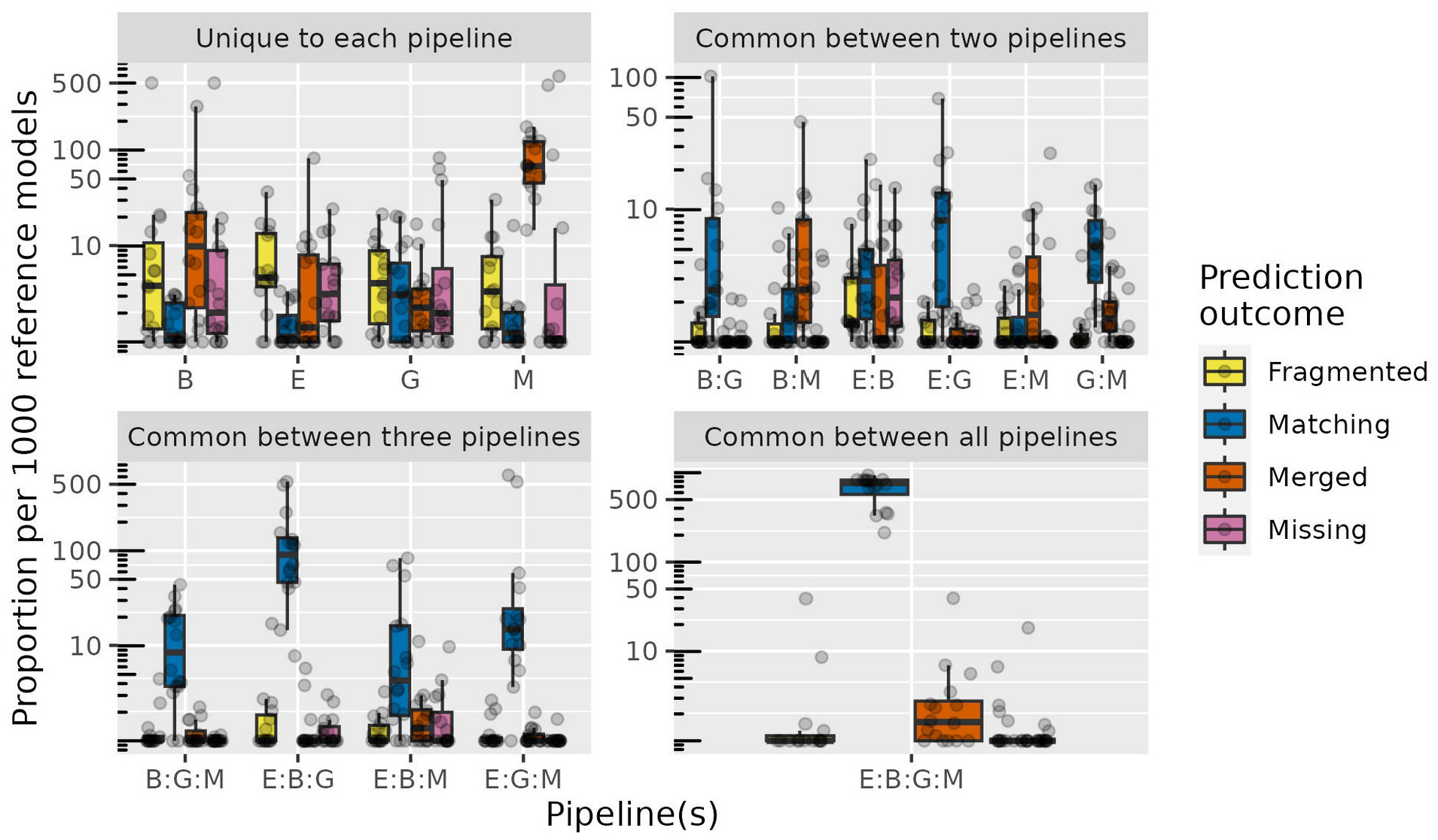
Boxplots (with corresponding data points overlayed) of proportions (per 1000 reference models) of gene prediction outcomes, per organism, common between two, three and four pipelines (e.g., E:G, number of gene predictions specifically made by Eukan and Gemoma but not the other two), as well as genes identified by one pipeline (E: Eukan; B: Braker; G: Gemoma; M: Maker). The trend is that gene predictions are generally correct and similar across three or four pipelines at a given locus, whereas incorrect predictions tend to unique to one or two pipelines only.

**Figure 5.**
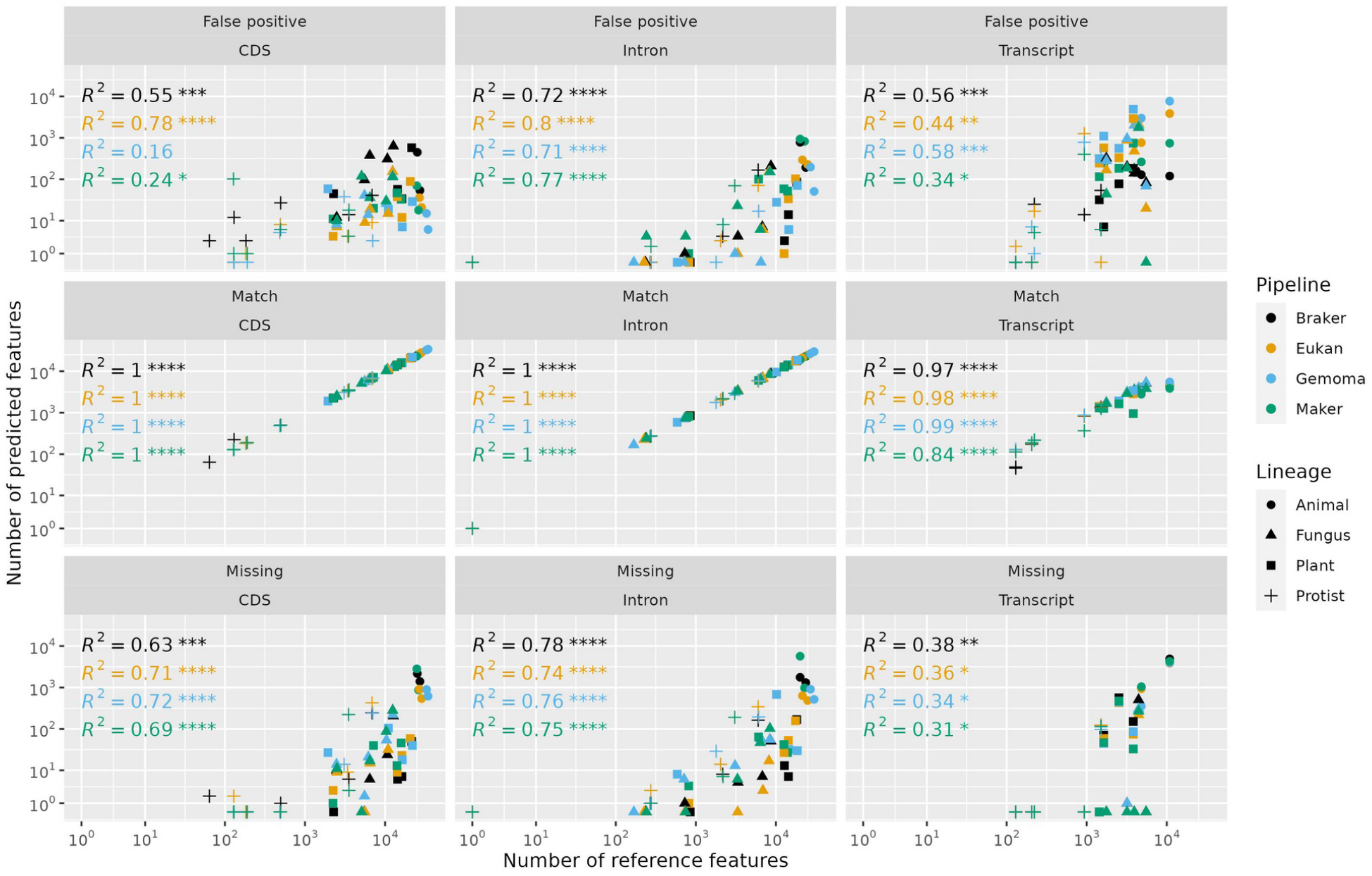
Scatterplots of the number of false positive, matching, and missing transcript, exon and intron predictions as a function of the number of corresponding reference features (curated, Busco-complete, or both) per annotation pipeline for each of the 17 tested genomes. The R^2^ values suggest a strong linear, and thus predictable, relationship (controlling for organisms without introns) between false positive, missing and matching predictions at the levels of transcripts, exons and introns (except false positive introns) for all pipelines. Moreover, there does not appear to be a significant difference in those trend lines. R^2^ represents the coefficient of determination, and the associated numbers of asterisks represent the level of significance (^*^ : p-value <0.01; ^**^ : p-value <0.001; ^***^ : p-value <0.0001).

Although the nuclear genomes used here for testing are taxonomically broader compared to previous studies, our results are still far from being representative for eukaryotes as a whole. Certain genomic features specific to some lineages frustrate attempts at correctly identifying coding regions, regardless of pipeline. For example, gene predictions in *Diplonema papillatum* (Diplonemea, Euglenozoa) informed by RNA-Seq data often end up with multiple distinct coding regions merged together (Valach, *et al*., 2023) because a considerable portion of genes are co-transcribed and then separated post-transcriptionally via spliced-leader *trans-*splicing (34). Theoretically, solely using protein sequence alignments to inform gene predictions is a solution but, in the case of *Diplonema*, would lead to equally undesirable outcomes. The large evolutionary distance between Diplonemea and the majority of studied species severely limits homology searches which provokes fragmented or missing predictions altogether.

Another difficult case are the *Blastocystis* species (Blastocystidae, Bigyra) which typically have compact genomes with short introns (median of 19 nt). Many protein-coding genes found within this lineage lack termination codons encoded in the genome but are added to the mRNA via a polyadenylation mechanism (Gentekaki et al., 2017). A combination of the former and latter provokes annotation pipelines to merge sometimes dozens of distinct coding regions. In a similar vein, RNA-editing (observed in diverse eukaryotic lineages (36)) reduces the utility of RNA-Seq-based evidence given the biological difference from the underlying genomic sequence. Mismatches are induced in read mapping which, in turn, skews coverage when taken as evidence, as well as hinders alignment of assembled transcripts.

Expansive genomes of some *Symbiodinium* species (Symbiodiniaceae, Dinophyceae), e.g., *S. kawaguti* (Lin et al., 2015), cause all existing annotation pipelines to fail spectacularly. The median exon length is 40 nt, spaced out by introns with a median length of 560 nt. Gene predictions are often truncated and inconsistent with the very evidence used to inform them (data not shown). Similarly, organisms with more complex chromosome architectures, such as polyploidy or heteroploidy, remain as much a challenge in annotation performance. Considerable tweaking of pipeline parameters, or manual intervention to devise case-specific workarounds may be required.

### False negatives and positives abound in modelling alternatively spliced transcripts

While pipelines generally perform well in identifying coding gene loci, transcript modelling yields highly variable results both between pipelines and for certain organisms. At the very least, an identified gene will contain one transcript prediction that matches a reference transcript. Predicted exons and introns therein will also be identified and modelled with high accuracy (Braker to a somewhat lesser extent, P<0.05). Yet, in most cases those correctly predicted transcripts can be among many other superfluously predicted transcripts, especially in organisms expected to express alternatives (particularly Gemoma, P<0.05). In contrast, false negatives are inevitable when modelling alternative splicing, at roughly at the same rate for all pipelines. While the latter appears to occur at a lesser rate, it arguably poses more of a challenge since absence of evidence can be caused by a number of factors outside the context of annotation.

Missing and superfluous transcript predictions by the tested pipelines are not surprising for three main reasons. First and foremost, expression of alternative transcripts depends on the physiological condition, developmental state, and for multicellular organisms, the tissue, which implies that a plethora of RNA-Seq data sets would need to be generated to more accurately reflect the transcriptome diversity. Even for well-studied model organisms these data are rarely available, which, incidentally, implies that the reference transcriptomes themselves are most likely deficient in various aspects. Second, Illumina reads (100-150 nt) used in this study are, on average, shorter than the transcript from which they were derived. Therefore, multiple variant transcripts resulting from alternative splicing events spaced further apart than the “short” read length lead to uncertainty in reconstructing accurate transcripts (38). The third reason relates to each pipeline’s alternative splicing algorithm, which, as shown in the Results section, generates considerably different outcomes given the same ‘Omics data.

The impact of false negative and false positive transcripts ripples down to the exon and intron predictions to varying degrees depending on the genome architecture. We observed that the numbers of missing and superfluous exon and introns depends on the intron density and known number of alternative transcripts. Organisms such as *D. melanogaster, C. elegans,* and *O. sativa* are on the high end of the spectrum, whereas intron-dense genomes with little-to-none (reported) alternative splicing in the reference, such as for *C. reinhardtii* and *T. gondii,* give rise to proportionally fewer false-negative and false-positive sub-features.

### Ongoing issues in annotation

We observed that Gemoma generated gene predictions whose cumulative F1 scores in three organisms (*C. primus, C. reinhardtii, D. discoideum*) deviated substantially from the other pipelines and, in addition, tended to generate significantly more false positive transcripts. In contrast, Maker was statistically more likely to merge adjacent coding gene regions, while Braker generated more false-positive exons in matching transcripts than the other pipelines, but the flip side is that it produced the fewest false-positive transcripts. Eukan appeared to perform consistently well in that it avoided generating extreme deviations compared to other pipelines. Parameters for each of the pipelines can be tuned to control their respective errors to some extent, yet no definitive framework exists to target a specific outcome without creating errors elsewhere.

Despite best efforts in parameterization, a small proportion of gene predictions were mis-identified (merged, split or missing) by the tested pipelines. One potential approach to mitigating this problem would be to first generate competing results from multiple pipelines in parallel; second, to identify conflicts in predicted gene intervals between those pipelines, and then search for an, e.g., three-out-of-four consensus. Such an algorithm would rectify a portion of defective models observed in this study, and possibly other organisms with similar genome architectures. In addition to being computationally intensive, the main drawback is that there is no procedure to assess correctness even if a majority of pipelines agree.

A more ideal solution would be to develop a novel algorithm that directly searches for protein sequences within the genome—one that is splice-site aware and employs Hidden Markov Models (HMMs) that are adequately representative of protein classes and as close to error-free as possible. The HMM itself could be used to assess prediction correctness, or to classify models similarly to Busco’s method.

False negative and, to a greater extent, false positive transcript models are an inherent feature of existing annotation pipelines. A promising approach towards more biologically relevant and complete modelling of transcript isoforms is full-length RNA sequencing via technologies such as ONT (39) and PacBio (40). A “long” read would, in theory, represent a full-length transcript and its splicing structures, thus uncertainty in attributing splice events to predicted transcripts can be largely avoided. Such long reads can also mitigate issues in gene mis-identification by providing more complete and contiguous information on expressed regions. However, a number of hurdles still exist (reviewed in (41). It is still cost prohibitive (in labour and resources) to catalogue a relatively complete transcriptome by combining data from multiple biological conditions. Some variant transcript structures will inevitably remain undocumented as we rarely know the conditions under which the diverse isoforms are expressed. Further, “long” sequence reads can also represent truncated transcripts, caused either by the RNA extraction procedure or from RNA degradation, which limits detection of variant structures the same way as in Illumina reads. Lastly, read quality is crucial to resolving transcript isoforms, as indels and mismatches within reads will cause mis-mapping and thus increase the probabilities of either missing or falsely predicting isoforms.

### Conclusion and Outlook

In this study, we presented a novel eukaryotic genome annotation pipeline, called Eukan, compared it to other actively maintained pipelines, and constructed a model for expected pipeline performance and expected error rates under certain conditions.

We demonstrated that Eukan generates high quality annotations more consistently with respect to those three other pipelines. The consistently high performance is due to its unique use of RNA-Seq read coverage to inform gene predictors and protein sequence alignments, along with its use of consensus gene algorithms optimized for general use. Another attractive feature of Eukan is that it comprises helper scripts to seamlessly run a functional annotation routine to prepare submission to public data repositories such as NCBI, as well as to generate an information-rich intron summary.

Annotation comparisons in the past were limited to a small set of curated models, but here we demonstrate how they can be supplemented with Busco assessments to provide unprecedented insight into annotation outcomes. An important result inferred from these augmented comparisons and from our novel pipeline is that, on average, the underlying quality of gene predictions are essentially indistinguishable between modern pipelines. This result, taken together with the fact that all modern pipelines build upon the same framework, would suggest that a radically different approach or breakthrough is necessary to make the next quantum leap in quality. This leap would ideally work towards reducing the types of gene prediction errors that continue to plague this generation of pipelines, as revealed by applying the classification system outlined here.

## Data availability

The code for Eukan is available on github at https://github.com/BFL-lab/eukan, and supplementary files available at https://megasun.bch.umontreal.ca/matt/eukan-manuscript. All accessions listed in the Supplementary Materials are available in the Genome and SRA databases of NCBI. Relevant code written for all analyses herein are available at https://megasun.bch.umontreal.ca/matt/.

## Key points

- We propose a novel eukaryotic genome annotation pipeline, called Eukan, and demonstrate its capabilities in consistently generating high-quality annotations.
- We describe a novel method for classifying the outcome of a gene predictions against a reference to create more actionable insights.
- We built a predictive model of performance for the tested pipelines which provides a crucial tool for framing expectations.
- We describe how the current generation of pipelines essentially converge in their performance and generate similar errors at similar rates, aside from a few exceptions.

