## Supplementary material for "Eukan: a fully automated nuclear genome annotation pipeline for less studied and divergent eukaryotes": Supplmentary Material

#### Methods

##### Genome assemblies

Eukaryotic organisms were selected where paired-end Illumina RNA-Seq data was available on SRA, genome assemblies and RNA-Seq data of those organisms were downloaded. The following fungal accessions were selected for download: *Aspergillus nidulans* (GCF_000149205.2), *Neurospora crassa* (GCF_000182925.2), *Saccharomyces cerevisiae* (GCA_000146045.2), *Schizosaccharomyces pombe* (GCF_000002945.1), *Ustilago maydis* (GCF_000328475.2). The following Protists accessions were selected: *Plasmodium falciparum* (GCF_000002765.3), *Trypanosoma brucei* (GCF_000210295.1), *Dictyostelium discoideum* (GCA_000004695.1), *Thalassiosira pseudoana* (GCA_000149405.2), *Leishmania major* (GCA_000002725.2), *Cyanidioschyzon merolae* (GCA_000091205.1) and *Toxoplasma gondii* (GCF_000006565.2). The two animal accessions were *Caenorhabdhitis elegans* (GCF_000002985.6) and *Drosophila melanogaster* (GCF_000001215.4). The selected plants were: *Arabidopsis thaliana* (TAIR10), *Oryza sativa* (GCA_001433935.1), *Chlamydomonas reinhardtii* (GCA_000002595.3), *Chloropicon primus* (GCA_007859695.1) and *Ostreococcus lucimarinus* (GCA_000092065.1)

##### Proteome selection

Ten proteomes of the closest phylogenetic neighbours were selected for each organism. *Aspergillus nidulans*: GCA_000002655.1, GCA_000002715.1, GCA_000002855.2, GCA_000006275.2, GCA_000149615.1, GCA_000149645.2, GCA_000184455.3, GCA_000239835.2, GCA_000600275.1, GCA_000812125.1. *Neurospora crassa*: GCA_000143365.1, GCA_000149955.2, GCA_000182805.2, GCA_000213175.1, GCA_000221225.1, GCA_000226095.1, GCA_000226115.1, GCA_000226545.1, GCA_001275765.2, GCA_900290415.1, GCF_000149205.2. *Saccharomyces cerevisiae*: GCA_000002515.1, GCA_000002525.1, GCA_000002545.2, GCA_000003835.1, GCA_000006335.3, GCA_000006445.2, GCA_000026365.1, GCA_000026945.1, GCA_001298625.1, GCA_001413975.1. *Schizosaccharomyces pombe*: GCF_000004155.1,

GCF_000149205.2, GCF_000149845.2, GCF_000149955.1, GCF_000150505.1, GCF_000182925.2, GCF_000349005.2, GCF_001477535.1, GCF_001477545.1, GCF_001661265.1. *Ustilago maydis*: GCA_000181695.1, GCA_000349305.2, GCA_000403515.1, GCA_000417875.1, GCA_000497045.1, GCA_000517465.1, GCA_000747765.1, GCA_003144125.1, GCA_900080155.1, GCA_900162835.1. *Plasmodium falciparum*: GCA_001680005.1, GCA_002157705.1, GCA_900002335.1, GCA_900005765.1, GCA_900005855.1, GCA_900090025.2, GCA_900090045.1, GCA_900097015.1, GCA_900240055.1. *Trypanosoma brucei*: GCA_000002725.2, GCA_000002845.2, GCA_000002875.2, GCA_000209065.1, GCA_000227375.1, GCA_000691245.1, GCA_001457755.2, GCA_002087225.1, GCA_003719475.1, GCA_003719485.1. *Dictyostelium discoideum*: GCA_000004695.1, GCA_000004825.1, GCA_000190715.1, GCA_000203815.1, GCA_000208925.2, GCA_000209125.2, GCA_000257125.1, GCA_000313135.1, GCA_000330505.1, GCA_000787575.2. *Thalassiosira pseudoana*: GCA_000296195.2, GCA_001750085.1, GCA_002217885.1, GCA_017506865.1, GCA_019154785.2, GCA_021029045.1, GCA_900660405.1, GCA_918797485.1, GCF_000149405.2, GCF_000150955.2. *Leishmania major*: GCA_001299535.1, GCA_009731335.1, GCA_017916305.1, GCA_017916325.1, GCA_017916335.1, GCA_017918235.1, GCF_000002845.2, GCF_000227135.1, GCF_000234665.1, GCF_001293395.1. *Cyanidioschyzon merolae*: GCA_000091205.1, GCA_000341285.1, GCA_000350225.2, GCA_002049455.2, GCA_003194525.1, GCA_008690995.1, GCA_013995675.1, GCF_000001735.4, GCF_000341285.1, GCF_000350225.1. *Toxoplasma gondii*: GCF_000006425.1, GCF_000006515.1, GCF_000165345.1, GCF_000208865.1, GCF_000223845.1, GCF_000499425.1, GCF_000499605.1, GCF_000499745.1, GCF_000769155.1, GCF_002563875.1. *Caenorhabdhitis elegans*: GCF_000001215.4, GCF_000002985.6, GCF_000005575.2, GCF_000149515.1, GCF_000183805.2, GCF_000371365.1, GCF_000507365.1, GCF_000956235.1, GCF_001040885.1, GCF_003254395.2. *Drosophila melanogaster*: GCF_000005215.3, GCF_000005925.1, GCF_000005975.2, GCF_000220665.1, GCF_000224195.1, GCF_000224235.1, GCF_000472105.1, GCF_000754195.2, GCF_002217835.1, GCF_003286155.1. *Arabidopsis thaliana*: GCF_000004255.2, GCF_000150535.2, GCF_000375325.1, GCF_000463585.1, GCF_000478725.1, GCF_000493195.1, GCF_000633955.1, GCF_000686985.2, GCF_000695525.1, GCF_000801105.1. *Oryza sativa*: GCF_000003195.3, GCF_000005005.2, GCF_000005505.3, GCF_000231095.2, GCF_000263155.2, GCF_001263595.1, GCF_001605985.2, GCF_002575655.2, GCF_016808335.1, GCF_018294505.1. *Chlamydomonas reinhardtii*: GCA_019650235.1, GCF_000001735.4, GCF_000092065.1, GCF_000143455.1, GCF_000147415.1, GCF_000214015.3, GCF_000258705.1, GCF_000611645.1, GCF_000733215.1, GCF_002220235.1. *Chloropicon primus*: GCF_000001735.4, GCF_000002595.1, GCF_000090985.2, GCF_000143455.1, GCF_000147415.1, GCF_000214015.3, GCF_000258705.1, GCF_000611645.1, GCF_000733215.1, GCF_002220235.1. *Ostreococcus lucimarinus*: GCF_000001735.4, GCF_000002595.1, GCF_000090985.2, GCF_000143455.1, GCF_000147415.1, GCF_000214015.3, GCF_000258705.1, GCF_000611645.1, GCF_000733215.1, GCF_002220235.1.

**RNA-Seq libraries**

The following Illumina RNA-Seq paired-end libraries were selected for each organism. *Aspergillus nidulans*: SRR4368892, SRR4368902, SRR4368910. *Neurospora crassa*:

SRR500048. *Saccharomyces cerevisiae*: SRR3396381, SRR3396382, SRR3396384, SRR3396385, SRR3396386, SRR3396387, SRR3396388, SRR3396389, SRR3396391, SRR3396392, SRR3396393. *Schizosaccharomyces pombe*: SRR097898, SRR097899, SRR097900, SRR097902, SRR097903, SRR097905, SRR097906, SRR097907, SRR097908, SRR097909, SRR097912, SRR097915, SRR097917, SRR097921, SRR097922, SRR097925, SRR402833. *Ustilago maydis*: SRR5235721. *Plasmodium falciparum*: SRR638979, SRR638980. *Trypanosoma brucei*: ERR141315. *Dictyostelium discoideum*: SRR6215636, SRR6215637, SRR6215638, SRR6215639, SRR6215640, SRR6215641, SRR6215642, SRR6215643, SRR6215644, SRR6215645, SRR6215646, SRR6215647, SRR6215648, SRR6215649, SRR6215650, SRR6215651, SRR6215652, SRR6215653, SRR6215654, SRR6215655, SRR6215656, SRR6215657, SRR6215658, SRR6215659, SRR6215660, SRR6215661, SRR6215662, SRR6215663, SRR6215664, SRR6215665, SRR6215666, SRR6215667, SRR6215668, SRR6215669, SRR6215670, SRR6215671, SRR6215672, SRR6215673, SRR6215674, SRR6215675, SRR6215676, SRR6215677, SRR6215678, SRR6215679, SRR6215680, SRR6215681, SRR6215682, SRR6215683, SRR6215684, SRR6215685. *Thalassiosira pseudoana*: SRR13953375, SRR13953376, SRR13953377, SRR13953378, SRR13953379, SRR13953380, SRR13953381, SRR13953382, SRR13953383, SRR13953384, SRR13953385, SRR13953386, SRR13953387, SRR13953388, SRR13953389, SRR13953390. *Leishmania major*: ERR2604475, ERR2604476, ERR2604477, ERR2604478, ERR2604479, ERR2604480. *Cyanidioschyzon merolae*: SRR16547541, SRR16547542. *Toxoplasma gondii*: SRR17053198, SRR17053199, SRR17053200. *Caenorhabdhitis elegans*: SRR065719. *Drosophila melanogaster*: SRR023505, SRR023546, SRR023608, SRR026433, SRR027108. *Arabidopsis thaliana*: SRR934391. *Oryza sativa*: SRR17779283, SRR17779284, SRR17779285, SRR17779286, SRR17779287, SRR17779288, SRR17779289, SRR17779290, SRR17779291, SRR17779292, SRR17779293, SRR17779294. *Chlamydomonas reinhardtii*: SRR15481341, SRR15481342, SRR15481343. *Chloropicon primus*: SRR8992761. *Ostreococcus lucimarinus*: SRR1300254. RNA-Seq reads were trimmed of any remaining adapter sequences with trimmomatic v0.35 [citation] and corrected with Rcorrector v1.0.4 [citation]. Corrected reads were mapped to their respective genome assemblies using STAR v2.7.3a [citation]. Intronic intervals were extracted from the STAR BAM file and formatted as GFF. Coverage was extracted in GFF format using bam2wig and wig2hints.pl bundled with Augustus v3.3.3 [citation]. Genome assemblies were initially masked using RepeatModeler v2.0.2a [citation]. A de novo transcriptome assembly of the RNA-Seq reads was done with Trinity v2.12.0 [citation]. Furthermore, a genome-guided transcriptome assembly was done with Trinity v2.12.0 using the STAR output BAM file. Pasa v2.4.1 [citation] was run on both the de novo and genome-guided transcriptome assemblies to create a comprehensive, non-redundant set of transcripts (following the developer guidelines at https://github.com/PASApipeline/PASApipeline/wiki). Exonic intervals from transcript alignments were extracted and formatted similarly to intron and coverage GFF files.

**Validated gene models.** The curated gene models for 12 of the here-tested organisms were retrieved from the 'Reviewed' Swiss-Prot sequence collection (accessed 2022-09-14). Sequences were selected where there was an explicit link in Swiss-Prot to a corresponding accession in the reference GFF3 files, and separately downloaded (from NCBI; accessions listed below) for the tested organisms. Sequences were further filtered to select only those supported by experimental evidence at either the transcript or the protein level. The number of records retrieved were *Arabidopsis thaliana:* 398; *Caenorhabdhitis elegans:* 331; *Chlamydomonas reinhardtii:* 28; *Cyanidioschyzon merolae:* 1; *Dictyostelium discoideum:* 1,264; *Drosophila melanogaster:*

4626; *Neurospora crassa:* 301; *Oryza sativa:* 2,348; *Plasmodium falciparum* 3D7: 55; *Saccharomyces cerevisiae* S288C: 5,503; *Schizosaccharomyces pombe:* 2483; and *Toxoplasma gondii:* 2.

**Preprocessing**

RNA-Seq reads were trimmed of remaining adapter sequences with Trimmomatic v0.35 (1) and corrected with Rcorrector v1.0.4 (2). Corrected reads were mapped to their respective genome assemblies using STAR v2.7.3a (3). Intronic coordinates were converted from STAR's splice junction output file to GFF using a custom script. Coverage was extracted using bam2wig and wig2hints.pl bundled with Augustus v3.3.3 and again formatted as GFF. Genome assemblies were masked using RepeatModeler v2.0.2a (4). The transcriptome was assembled *de novo* from RNA-Seq reads with Trinity v2.12.0 (5). Furthermore, a genome-guided transcriptome assembly was done with Trinity v2.12.0 using the STAR-produced BAM file. Pasa v2.4.1 was run on both the *de novo* and genome-guided transcriptome assemblies to create a comprehensive, non-redundant set of transcripts (following the developer guidelines at https://github.com/PASApipeline/PASApipeline/wiki). Exonic coordinates from transcript alignments were extracted and formatted similarly to the intron and coverage GFF files mentioned above.

**Design principles behind Eukan**

We sought to design a pipeline that leveraged a broader range of information, with the expectation that the gene modelling process will be improved by incorporating under-exploited forms of evidence, notably RNA-Seq read coverage and intronic genome regions inferred from split reads. In addition, we aimed at using conventional evidence in novel ways by, for example, informing protein sequence alignments with expected intron length distributions, informing gene predictions with RNA-Seq coverage data, and using an optimized evidence weighting scheme of gene predictions, protein and transcript sequence alignments (empirically-derived from testing various weight combinations to yield the best over all increase in F1 scores; data not shown) for building consensus models. Finally, unlike traditional pipelines, ours include auxiliary scripts that will also associate functional information to gene models, as well as to summarise intron statistics. The pipeline, called 'Eukan', has been used successfully in the meantime in several nuclear genomes studies of eukaryotes from most diverse groups such as jakobids, amoebozoa, leotiomycetales fungi, diplonemids (Gray *et al.*, 2020; Matthey-Doret *et al.*, 2022; Thimmappa *et al.*, 2023; Valach *et al.*, 2023), and dinoflagellates (unpublished). The challenges particular to each genome allowed us to incrementally improve the pipeline to its current state.

For benchmarking, we compare the performance of Eukan with recent versions of the three above described annotation pipelines on a broad range of eukaryotic genomes (animals, fungi, plants and protists). For that, we developed a classification scheme (abstracted layer) for the standard sensitivity and specificity originally defined by Burset and Guigo (1996) and elaborated in the Eval package (9).

Further, for a more comprehensive performance test, we passed the pipelines not as usually done just through the genomes from a few animals, fungi, and plants, but on a total of 17 genome assemblies with available annotations from phylogenetically diverse taxa, notably *Plasmodium falciparum* (Alveolata), *Dictyostelium discoideum* (Amoebozoa), *Trypanosoma brucei* and *Leishmania major* (Kinetoplastida), *Cyanidioschyzon merolae* (Rhodophyta), *Toxoplasma gondii* (Apicomplexa); the ascomycete fungi *Aspergillus nidulans, Neurospora crassa,* and *Saccharomyces cerevisiae, Schizosaccharomyces pombe,* and the basidiomycete d gene models correspond to 'complete' Busco*Ustilago maydis; Chlamydomonas reinhardtii*

(Chlorophyta), and the streptophyta *Chloropicon primus*, *Oryza sativa*, along with the model plant *Arabidopsis thaliana*; and two model animals, *Caenorhabdhitis elegans* and *Drosophila melanogaster*, the latter three being commonly used for performance testing in publications on annotation pipelines.

**Executing the pipelines**

To run Braker, the following input files were specified: the assembly of the genome of interest, the hints file containing intronic regions (and their respective coverage) in GFF format, and the proteome fasta files from ten related species. Additional options invoked are '--softmasking', '--etpmode', '--alternatives-from-evidence' and, when applicable, '--fungus'. The input files provided to Gemoma include the genome assembly, the assemblies of neighbouring organisms and their corresponding annotations in gff format, and the bam file generated by Star. The 'CLI GeMoMaPipeline' method was used, specifying 'GeMoMa.Score=ReAlign', 'AnnotationFinalizer.r=NO', 'AnnotationFinalizer.u=YES' and 'r=MAPPED'. The Maker pipeline was executed similarly as described earlier (10) and recommended by the developers of Maker, but with several modifications to increase the confidence in hints derived from sequence alignments. Specifically, 'e*_score_limit' was changed from 20 to 30, 'pcov_blastn' was changed from 0.8 to 0.9, 'pid_blastn' was changed from 0.85 to 0.95, 'min_contig' was changed from 10kb to 4kb, 'single_exon' was changed from 0 to 1, 'keep_preds' was changed from 0 to 1. The option 'alt_splice' was set to 1 at the final step of the pipeline to identify alternative transcripts. The Augustus optimization step was also run to better replicate the steps taken by Eukan and Braker. The Eukan script was launched with the genome assembly file, proteome sequence files, a transcript sequence fasta and gff3, and hints file in GFF format (specifying 'exon', 'exonpart' and 'intron' hints; see Eukan workflow Section). Additional boolean flags used are '--strand_specific_transcripts', '--utrs' and, where applicable, `--fungus`. The exact commands to run each pipeline on each organism are further detailed in the Supplementary Material.

**Gene prediction quality statistics**

The prediction quality metrics implemented in this study are Sensitivity (Sn), Precision (Pr) and harmonic mean (F1), as defined by others (11), applied at the level of genes, transcripts, exons and introns, and as formalized in, e.g., the Eval package (9). Briefly, these statistics compare the coordinate overlap between two features of the same type (e.g., the coordinates of the predicted and the reference exon) at a given genomic locus. Sn is the fraction of a reference model locus that overlaps with a prediction. In other words, it is the true positive (TP) length divided by the sum of the TP and false negative (FN) portions of the overlap, $Sn=TP/(TP+FN)$. Pr is defined as the fraction of a predicted locus that overlaps with a reference locus, i. e., the TP length divided by the sum of the TP and false positive (FP) segments, $Pr=TP/(TP+FP)$. The harmonic mean is computed by the following formula: $F1=2TP/(2TP+FP+FN)=2Sn\times Pr/(Sn+Pr)$. The statistics range from 0 (no overlap) to 1 (full overlap). F1 is sometimes interchangeably referred to as Accuracy (Acc), and Precision is sometimes interchangeably referred to as Specificity (Sp) (10, 12).

**Statistical analyses implemented in comparisons**

Statistical analyses were carried out using R v4.1.2. To enhance resolution, comparisons of empirical cumulative distribution functions (ECDFs) (of F1 scores and Busco results) were carried out using the DTS test in the 'two-samples' package (v2.0.0, Dowd, 2020). Correlation coefficients and coefficients of determination ($R^2$, with P-values calculated by the Spearman method (14) were computed for prediction outcome categories (e.g., 'matching', 'missing', ‘fragmented’, ‘FP’) using the ggpubr package (v0.6.0, Kassambara and Kassambara, 2020). Linear models were fitted to those prediction outcomes using lstrends from the emmeans package (v1.8.5, Lenth et al., 2019). Here, trend lines (obtained for each pipeline and prediction feature gene, transcript, exon, intron) were compared using the pairs test from the ‘graphics’ package (v3.6.5, Murrell, 2005). Multiple test corrections were carried out using the ‘BH’ method (18) implemented in the R ‘stats’ package (v4.3.0, Team et al., 2018). Non-parametric post-hoc tests (20) were computed using the FSA R package (v0.9.4, Ogle and Ogle, 2017).Missing, split and merged predictions were generated by all p

### Supplementary Results

##### Assessment of pipeline-predicted genes on the extended gold standard collections

The quality of gene predictions made by the tested pipelines is discussed here, first on the Busco assessment of conceptually translated protein-coding genes, then through the classification framework defined in the Methods section.

The conceptually translated sequences of protein-coding gene predictions across all tested organisms by Braker, Eukan, Genoma and Maker were compared using their respective assessments by Busco, i.e., proportions of ‘complete’, ‘missing’ and ‘fragmented’. Cumulatively, more than 75% of pipeline-predicted gene models correspond to ‘complete’ Buscos (Supplementary Figure 1). Exceptionally high rates of ‘missing’ Busco models were observed in Maker runs on *O. sativa* and *T. gondii* (>60%), and Braker on *T. Brucei* (>40%). In contrast, gene predictions deemed ‘fragmented’ by Busco occurred at rates even lower than those that are missing (median ~0.8%, mean ~1.6%) where fewer extreme cases were observed.

We classified the loci of predicted genes by the pipelines as either ‘matching’, ‘missing’, ‘fragmented’, or ‘merged’ (defined in Figure 2). All pipelines show a strong correlation between gene predictions that match their corresponding reference locus one-to-one (Figure 1). Most (>75%) of those gene predictions are exact matches to the 5` and 3` reference coordinates (median F1 of 1, mean F1 ~1, Supplementary Figure 4). Furthermore, all pipelines tended to predict genes at the same genomic locations (Figure 2), with no statistically apparent difference. That said, outliers in F1 distributions of matching models (i.e., more median F1 of 1, mean F1 ~1 imperfect models) were specifically observed for Gemoma in *C. primus, C. reinhardtii, D. discoideum,* Maker in *O. sativa, L. major*, in contrast to Eukan, which did not exhibit any extreme behavior.

Braker and Gemoma showed weak correlations in expected proportions of merged, missing and split models, while Maker exhibited a clear bias ($R^2$=0.84, P<0.0001) towards merging adjacent coding gene loci together where the reference (and other pipelines) suggests otherwise. Missing, split and merged predictions were generated by all pipelines to some extent (Figure 2), but erroneous predictions were more likely to be specific to the pipeline as opposed to a given genomic locus (P<0.05). An average of ~2.5% and ~3.5% of predictions at loci with Busco-complete reference models were found to be split and merged, respectively.

##### Assessment of pipeline-predicted transcripts on the extended gold standard collections

The proportions of single-copy, duplicated, missing and fragmented transcript predictions per pipeline, according to Busco assessments, are summarized in Supplementary Figure 3. Large discrepancies in proportions of duplicated transcript sequences were observed between pipelines for organisms known to have elevated rates of alternative splicing (e.g., *C. elegans, D. melanogaster, O. sativa*). Surprisingly, large discrepancies in rates of predicted isoforms were also observed between pipelines in certain organisms where few cases of alternative splicing have been documented (e.g., *A. nidulans*, *S. pombe*). Nevertheless, all the tested pipelines tended to predict at least one transcript that matched a corresponding reference transcript (Figure 3). Furthermore, the F1 scores of those predicted transcripts were generally high, e.g., >75% of predictions were exact matches to their corresponding reference (Supplementary Figure 4). No significant difference in F1 scores was observed between pipelines yet, similarly to the gene predictions, some outliers were observed in certain cases. Gemoma predicted considerably more imperfect models (>10%) in *C. primus* , *C. reinhardtii*, and *D. discoideum*. Maker generated more imperfect models than the other pipelines on *O. sativa* and *L. major* by about 10%. Braker generated about 10% more imperfect models in *S. pombe* compared to other pipelines. Eukan was the only pipeline that did not suffer any outlier outcomes in F1 scores of matching predictions in any of the tested organisms.

**Assessment of pipeline-predicted exons and introns on the extended gold standard collections**

Exons and introns were extracted strictly from predicted transcripts that correspond (perfectly or imperfectly) with a reference transcript, i.e., features of false positive transcripts were ignored. All pipelines succeeded in identifying the majority (mean >95%) of reference exons (Supplementary Figure 4). The relatively high and statistically significant $R^2$ values observed for false negative exons suggest that a mean of about 5% will be missing in any given annotation run. Those missing exon loci account for most of the F1 score deviations observed in imperfectly matching transcript models (<25%). In contrast, false-positive exon predictions appear to be an order of magnitude less frequent than those of false negatives. Yet, Braker was found to significantly generate more false positive exons than other pipelines (P<0.05 Dunn post-hoc test, BH multiple test correction; no significant difference between the other three pipelines). Of all the pipelines, Eukan ($R^2$=0.79, P<0.0001) appears to more predictably generate a low rate of false positive exons compared to Braker ($R^2$=0.55, P<0.0001), Gemoma ($R^2$=0.15, ns) and Maker ($R^2$=0.24, P<0.05).

Across all pipelines, the degree of accuracy among the matching exon predictions was high (>95%, Supplementary Figure 4), with the exception of a few organism- and pipeline-specific outliers. Braker predicted around 8% more imperfect models than other pipelines in *S. pombe* and *C. merola*. Gemoma generated about 8% more imperfect exons than other pipelines in *C. primus*. Maker generated about 40% more imperfect exons in *L. major*. Eukan was not observed to generate outlier outcomes at the exon level. That said, internal exons were consistently predicted more accurately than 5`, 3` or single exons by all pipelines (Supplementary figure 6). The 5` exon start tended to be most variable, which can be explained by the presence of alternative in-frame start codons.

Consistent with the high rate of correctly identified exons, all pipelines were able to identify the vast majority of reference introns (>95%, Figure 1). The number of missing introns is consistently between 1%-5%, with no significant difference between pipelines. Incidentally, a missing intron prediction typically implies a corresponding missing exon, but a missing exon does not necessarily imply a missing intron due to single-exon genes. On the other hand, a false-positive intron is typically associated with a false-positive exon. The rate at which pipelines generated spurious introns was in the same order of magnitude as omitted introns (no significant difference), and at similar $R^2$ values.

More than >95% introns were correctly identified and demarcated, which is consistent with the F1 scores observed in internal exon predictions (Supplementary Figure 6). Thus, internal gene structure predictions were highly accurate. No pipeline- or organism-specific outliers were observed in F1 distributions like those observed at the gene, transcript, or CDS level.

#### Supplementary Tables

*Supplementary Table 1 : Breakdown of the numbers of reference models used as benchmarks for each tested organism. The datasets are comprised of both known curated models as well as the extended models identified as 'Complete' by Busco. There is considerable overlap between curated models and Busco hits in some cases, whereas 'complete' Busco hits in other cases provide substantial benchmark models.*

| Organism | Curated, non-Busco | Uncurated Busco complete | Curated & complete Busco overlap | Total |
|---|---|---|---|---|
| *S. cerevisiae S288C* | 3338 | 33 | 2144 | 5515 |
| *O. sativa* | 2142 | 1237 | 112 | 3491 |
| *S. pombe* | 1727 | 692 | 755 | 3174 |
| *D. melanogaster* | 1489 | 2612 | 587 | 4688 |
| *D. discoideum* | 1120 | 227 | 25 | 1372 |
| *A. thaliana* | 266 | 1598 | 22 | 1886 |
| *C. elegans* | 232 | 3090 | 26 | 3348 |
| *N. crassa* | 98 | 3634 | 161 | 3893 |
| *P. falciparum 3D7* | 51 | 167 | 4 | 222 |
| *C. reinhardtii* | 22 | 1514 | 1 | 1537 |
| *T. gondii* | 2 | 894 | 0 | 896 |
| *C. merolae* | 1 | 188 | 0 | 189 |
| *A. nidulans FGSCA4* | 0 | 3808 | 0 | 3808 |
| *C. primus* | 0 | 1428 | 0 | 1428 |
| *L. major* | 0 | 130 | 0 | 130 |
| *T. brucei* | 0 | 128 | 0 | 128 |
| *U. maydis 521* | 0 | 1755 | 0 | 1755 |

### Supplementary Figures

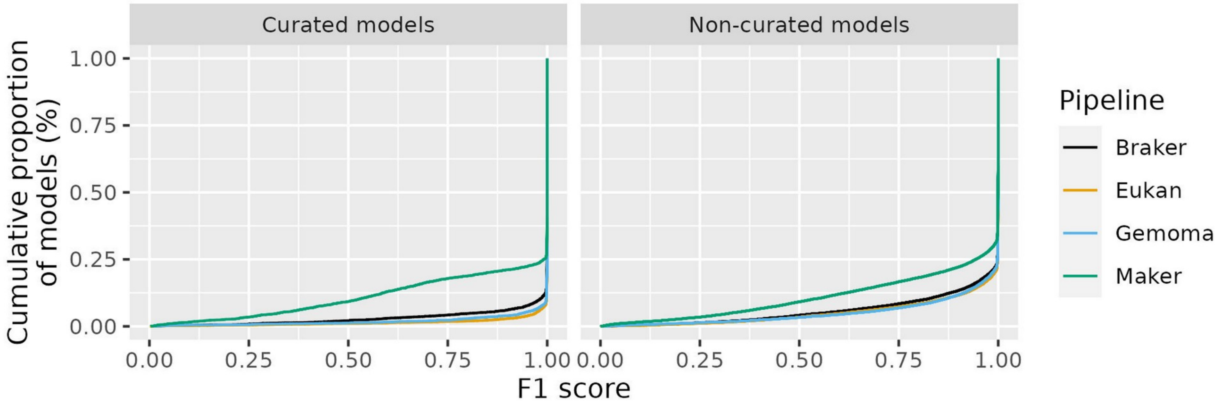


*SupplementaryFigure 1: Cumulative F1 scores of gene predictions, generated by each pipeline and for all tested organisms, at genomic loci of known curated reference models (left) and non-curated reference models found to be 'complete' by Busco.*

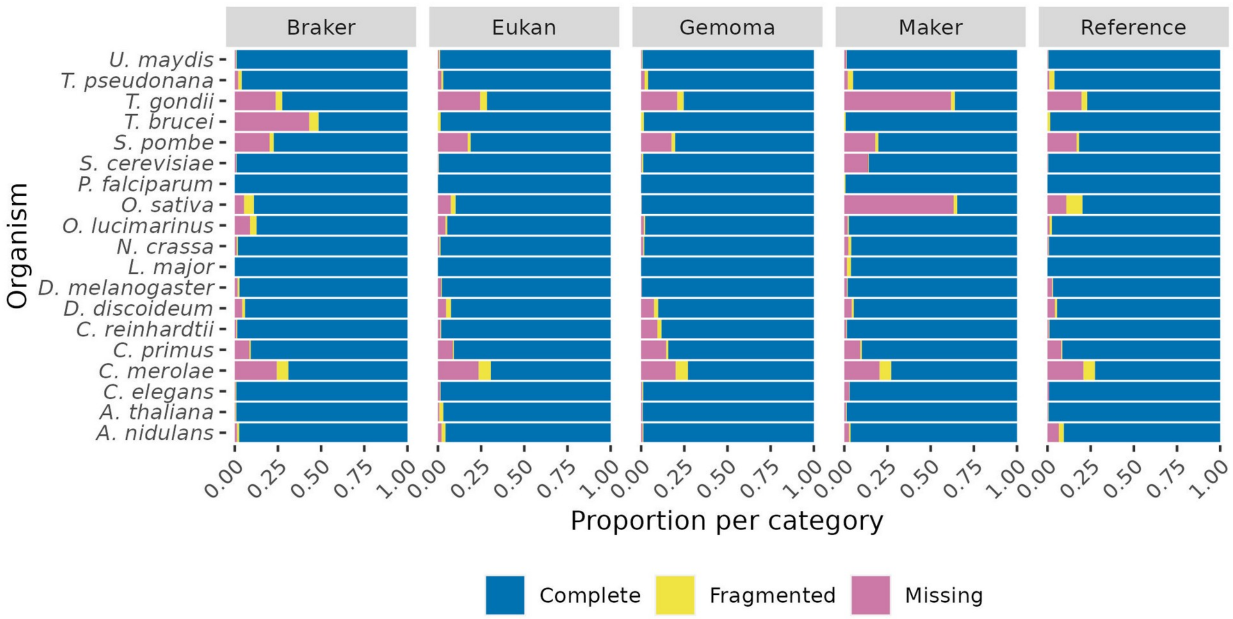


*SupplementaryFigure 2: Stacked barchart of proportions reported in 'Busco completeness assessments' for (conceptually translated) genes in the the reference set, as well as those generated by Braker, Eukan, Gemoma and Maker on the 17 tested genomes. Proportions correspond to the relative numbers of gene models identified by Busco within the respective lineage-specific OrthoDBv10 dataset. Proportions of complete vs fragmented vs missing genes are generally consistent across pipelines and reference per organism, aside from some visible differences in T. gondii, T. brucei, O. sativa, C. reinhardtii.*

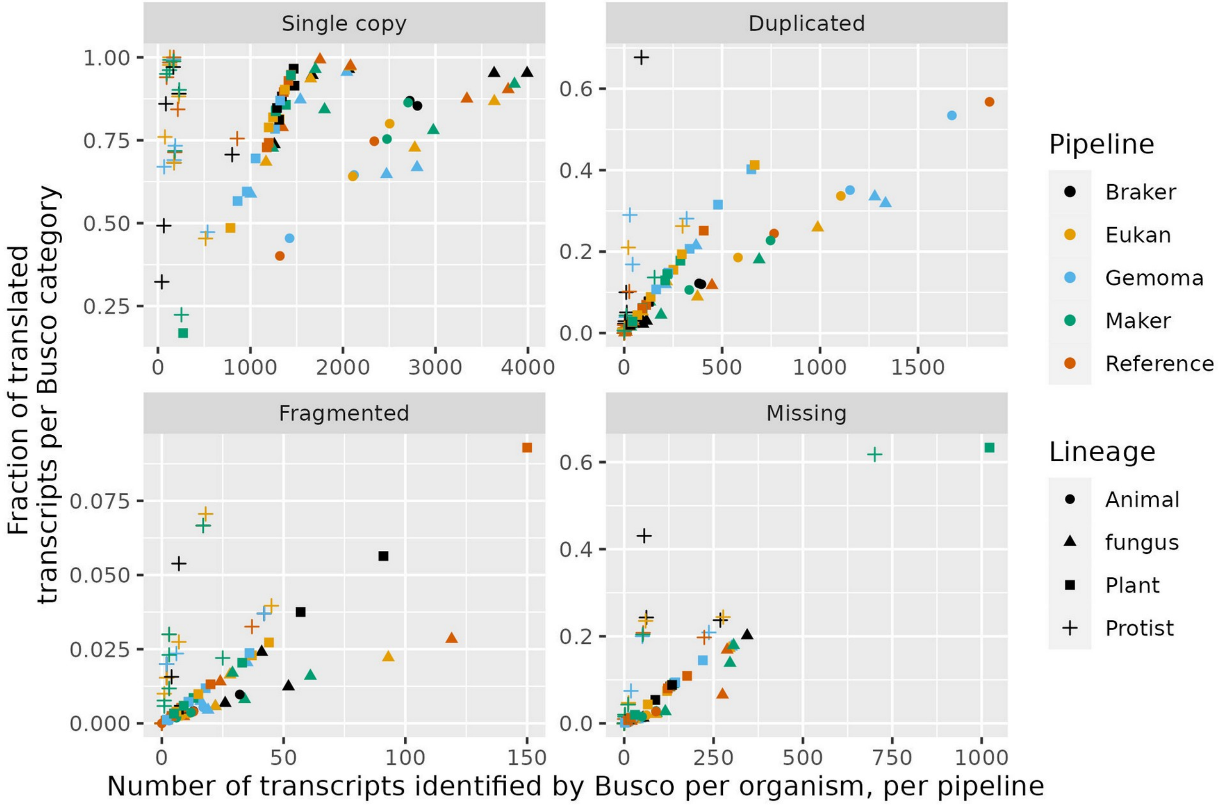


*Supplementary Figure 3: Scatterplots of (conceptually translated) transcripts identified as either single copy, duplicated, fragmented or missing by Busco for both the reference transcripts and predicted transcripts (by all pipelines) for the 17 tested organisms. Numbers along the horizontal axis correspond to the absolute numbers identified by Busco from the respective lineage-specific OrthoDBv10 dataset. The vertical axis corresponds to the percentage breakdown as reported in Busco assessments.*

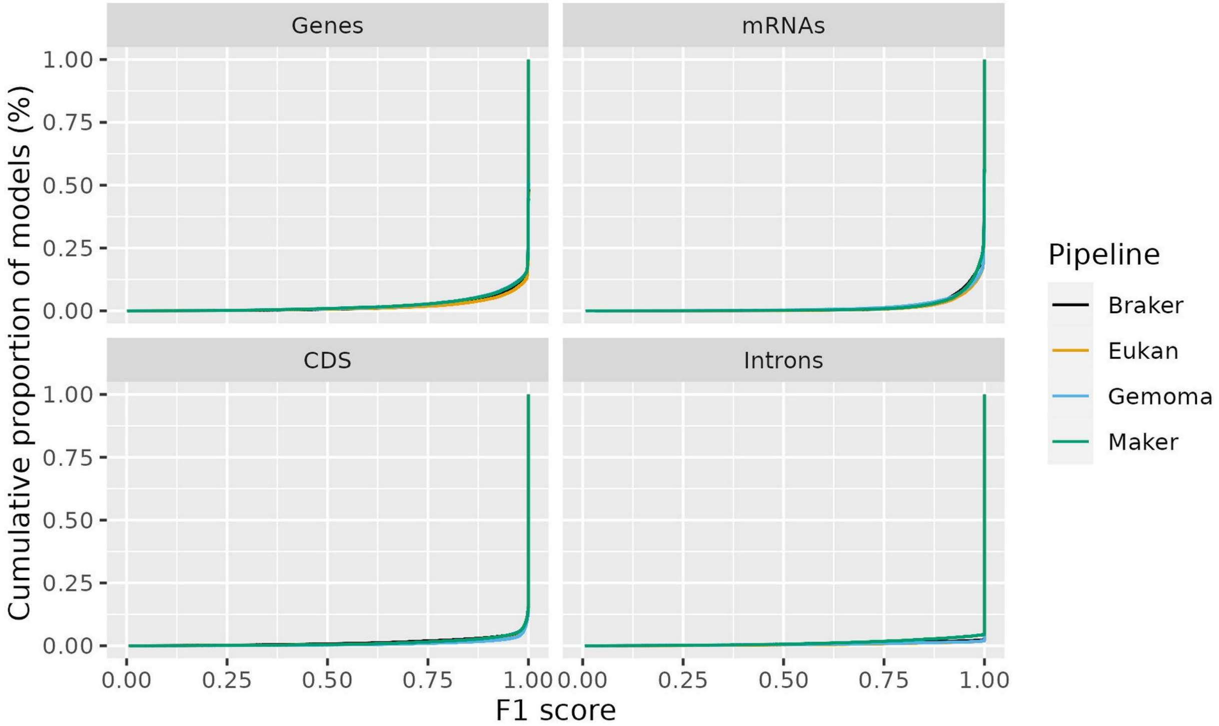


*SupplementaryFigure 4: Cumulative F1 scores for all gene, mRNA, CDS and intron predictions by each pipeline across the tested reference annotations.*

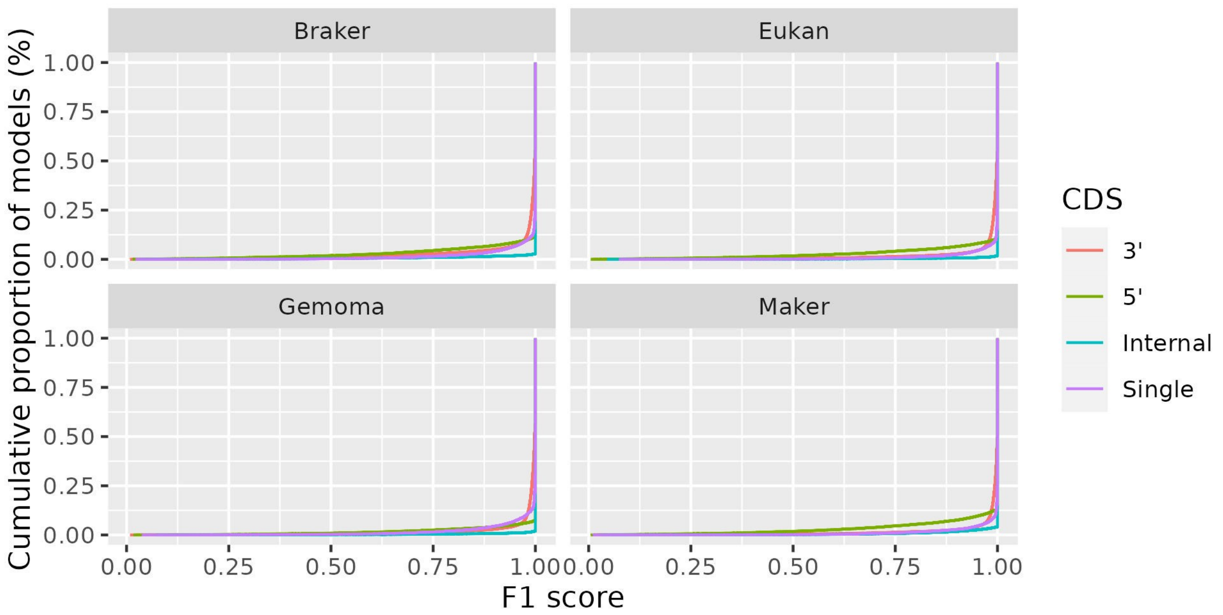


*SupplementaryFigure 5: Cumulative F1 distributions of initial, internal, terminal and single-exon features predicted by each pipeline. The distributions in all cases are skewed towards an F1 score of 1, wherein internal CDS scores are most skewed, suggesting extremely high prediction accuracy.*

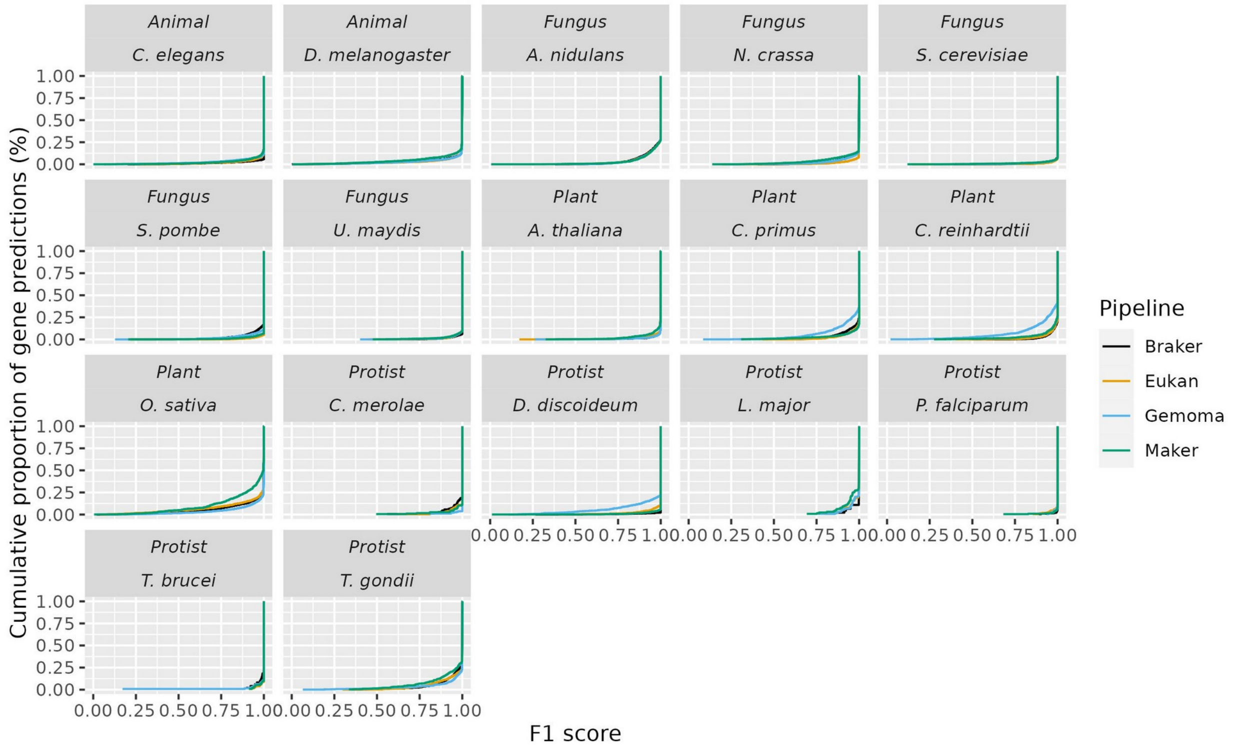


*SupplementaryFigure 6: Empirical cumulative distributions of gene prediction F1 scores that 'match' a corresponding reference, generated by the four pipelines grouped per organism. Gene predictions made by all pipelines match the reference exactly 75%, except C. primus, C. reinhardtii, O. sativa, D. Discoideum where one pipeline deviated considerably from the other three.*

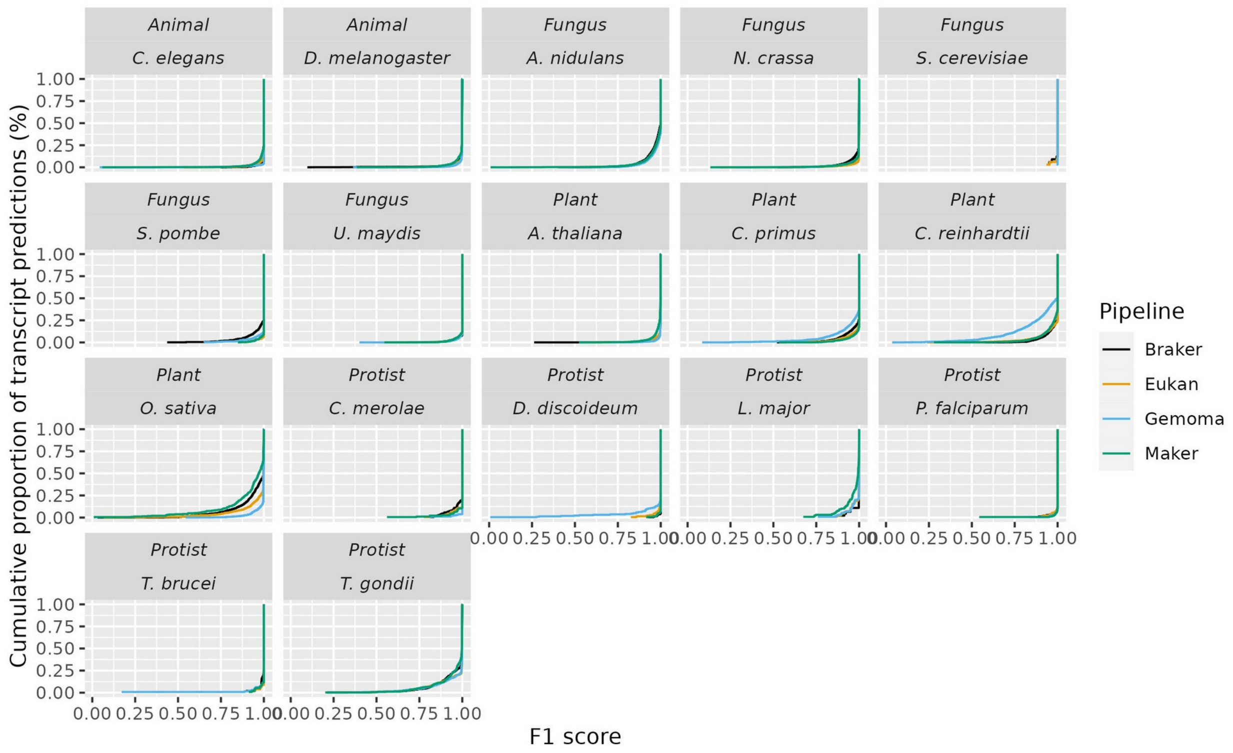


*SupplementaryFigure 7: Empirical cumulative distributions of transcript prediction F1 scores that 'match' a corresponding reference, generated by the four pipelines grouped per organism. Transcript predictions made by all pipelines with a corresponding match in the reference were generally exact (>75%). Exceptions to this trend were observed in 1) A. nidulans where all pipelines predicted ~50% of the corresponding references exactly, 2) Gemoma transcript predictions in C. reinhardtii and L. major, and 3) the spread in pipeline performance in O. sativa.*

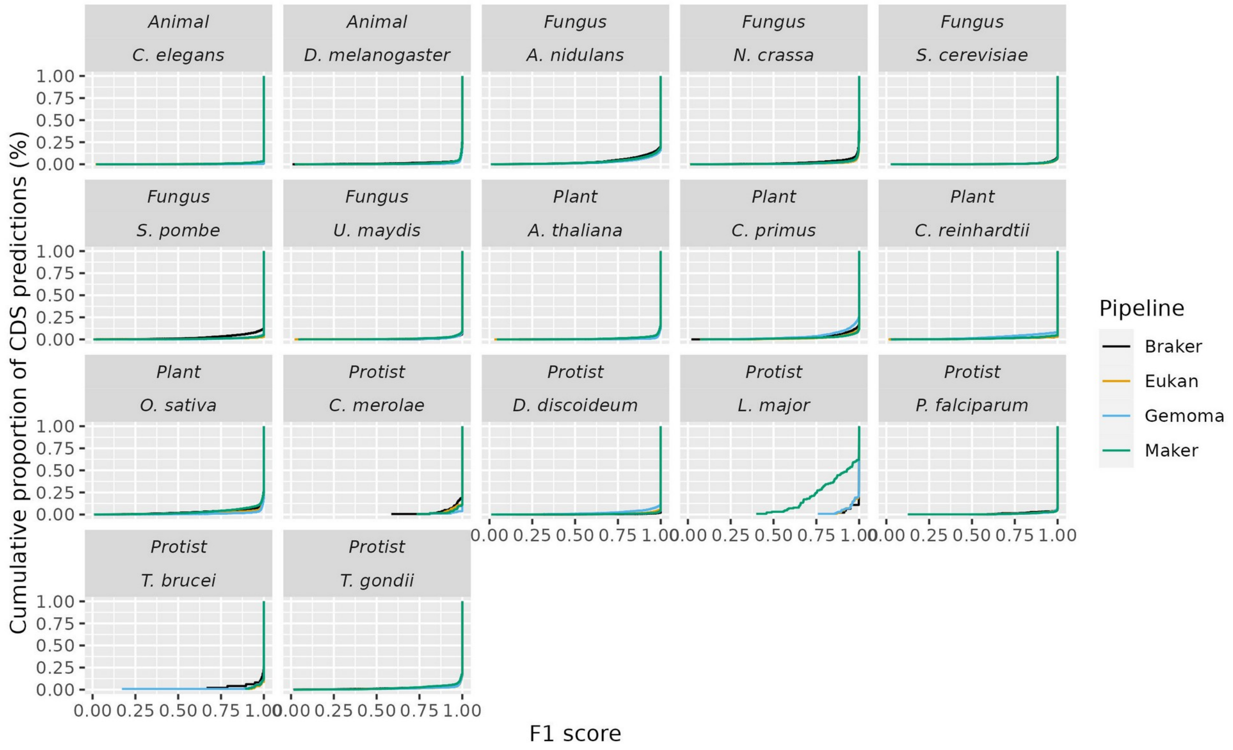


*SupplementaryFigure 8: Empirical cumulative distributions of exon prediction F1 scores that 'match' a corresponding reference, generated by the four pipelines grouped per organism. CDS prediction quality tended to be high (>80% exact matches) and consistent between pipelines, save for Maker predictions in L. major.*

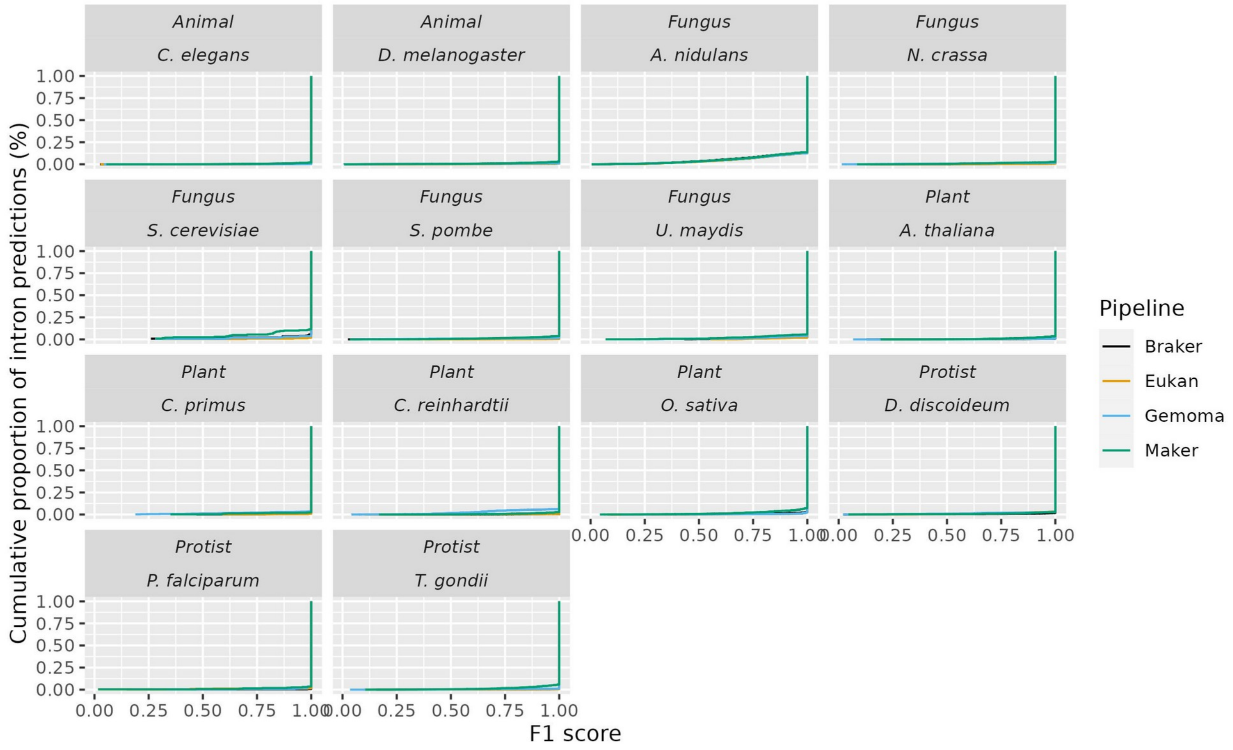


*SupplementaryFigure 9: Empirical cumulative distributions of intron prediction F1 scores that 'match' a corresponding reference, generated by the four pipelines grouped per organism. Intron prediction quality tended to be high (>80% exact matches) and consistent between pipelines.*
